# Glyoxalase 1: emerging biomarker and therapeutic target in cervical cancer progression

**DOI:** 10.1101/2024.02.11.579832

**Authors:** Ji-Young Kim, Ji-Hye Jung, Soryung Jung, Sanghyuk Lee, Hyang Ah Lee, Yung-Taek Ouh, Seok-Ho Hong

## Abstract

**Introduction:** Cervical cancer presents a significant global health challenge, disproportionately impacting underserved populations with limited access to healthcare. Early detection and effective management are vital in addressing this public health concern. This study focuses on Glyoxalase-1 (GLO1), an enzyme crucial for methylglyoxal detoxification, in the context of cervical cancer.

**Methods:** We assessed GLO1 expression in cervical cancer patient samples using immunohistochemistry. *In vitro* experiments using HeLa cell lines were conducted to evaluate the impact of GLO1 inhibition on cell viability and migration. Single-cell RNA sequencing (scRNA-seq) and gene set variation analysis were utilized to investigate *GLO1*’s role in the metabolism of cervical cancer. Additionally, public microarray data were analyzed to determine *GLO1* expression across various stages of cervical cancer.

**Results:** Our analysis included 58 cervical cancer patients, and showed that GLO1 is significantly upregulated in cervical cancer tissues compared to normal cervical tissues, independent of pathological findings and disease stage. *In vitro* experiments indicated that *GLO1* downregulation decreased cell viability and migration in cervical cancer cell lines. Analyses of scRNA-seq data and public gene expression datasets corroborated the overexpression of *GLO1* and its involvement in cancer metabolism, particularly glycolysis. An examination of expression data from precancerous lesions revealed a progressive increase in *GLO1* expression from normal tissue to invasive cervical cancer.

**Conclusions:** This study highlights the critical role of GLO1 in the progression of cervical cancer, presenting it as a potential biomarker and therapeutic target. These findings contribute valuable insights towards personalized treatment approaches and augment the ongoing efforts to combat cervical cancer. Further research is necessary to comprehensively explore GLO1’s potential in clinical applications.

**Synopsis:** This study examines Glyoxalase-1 (GLO1) in cervical cancer and reveals its significant upregulation in cancerous tissues compared to normal counterparts. *In vitro* experiments show that downregulating GLO1 decreases cell viability and migration. Single-cell RNA sequencing and gene expression analysis substantiate GLO1’s involvement in cancer metabolism, notably in glycolysis. These findings identify GLO1 as a potential biomarker and therapeutic target, offering insights for personalized treatment approaches in cervical cancer management.

## Introduction

Cervical cancer, a prevalent and overwhelming malignancy among women, presents a significant global health challenge [1]. Its severity stems not only from high incidence and mortality rates but also from the disproportionate burden it imposes on underserved populations with limited healthcare access [2, 3]. Timely detection and effective management of cervical cancer are crucial, and identifying novel biomarkers is a key step in addressing this urgent public health issue [2]. Understanding the complex pathogenesis of cervical cancer remains a multifaceted challenge [4]. Although the link between human papillomavirus (HPV) infection and cervical cancer is well-established, the disease’s pathogenesis involves various molecular and cellular mechanisms that are not yet fully elucidated [5]. Additionally, cervical cancer displays significant heterogeneity in progression and behavior, with some cases being more aggressive than others [6]. This variability highlights the need for comprehensive insights into the underlying pathogenic processes, including the identification of genetic, epigenetic, and environmental factors contributing to disease development. A deeper understanding of these pathways is essential for developing targeted prevention and treatment strategies, aiming to reduce the global burden of cervical cancer.

Glyoxalase-1 (Glo1) has emerged as a key player in cancer progression and prognosis. Aberrant expression and activity of Glo1 have been linked to the development, aggressiveness, and treatment resistance of various cancer types. Published studies have shown that the expression of Glo1 is upregulated in breast cancer tissues, and that patients with stage 3-4 tumors with high GLO1 and PKCλ expression have significantly poor survival compared to patients with low expression of these genes. Treatment of MDA-MB-157 and MDA-MB-468 human basal-like breast cancer cells with TLSC702, a GLO1 inhibitor, resulted in decreased cancer cell survival and tumor-sphere formation, suggesting that GLO1 is involved in cancer progression and contributes to the poor prognosis of breast cancer [7, 8]. GLO1 plays a major role is to catalyze the conversion of glycolytic methylglyoxal (MG) to non-toxic D-lactate, and it represents a tumor survival strategy by inhibiting the accumulation of cytotoxic MG in highly metabolized tumor tissues. In particular, metastatic prostate cancer patients have higher levels of GLO1 and significantly lower levels of MG compared to non-metastatic prostate cancer patients, and GLO1 siRNA-mediated depletion in PCa cell lines induces the accumulation of MG-derived AGEs and further inhibits the expression of EMT-related genes such as E-cad and ZO-1, which are involved in cancer cell migration and invasion. These findings indicate that GLO1 promotes survival by suppressing intracellular levels of the cytotoxic MG-derived AGEs, thereby avoiding cell death [9–11]. Upregulated expression of Glo1 is associated with multi-drug resistance(MDR) leading to chemotherapy failure, which in turn is linked to cancer cell survival and development. It has been reported that GLO1 is increased 2.4-fold in Ishikawa cells, an endometrial cancer cell line, compared to parental cells, and is associated with progestin resistance in endometrial carcinoma. Focusing on GLO1 as a mediator of progestin resistance in endometrial cancer patients, and metformin treatment of endometrial cancer cells alleviated progestin resistance and reduced cell proliferation through dysregulation of GLO1 [12]. In metastatic prostate cancer, overexpression of GLO1 induced MG-H1-mediated upregulation of PD-L1 to promote cancer progression, and increased expression of GLO1 in clinical chemotherapy of breast cancer reduces the cytotoxicity of anticancer drugs, contributing to MDR [13, 14]. In another study, cancer stem cells (CSCs) are known to influence MDR and recurrence, and Glo1 was reported to be essential for CSC growth and survival. In breast cancer, CSCs are characterized by a CD44hi/CD24-/low phenotype and expression of ALDH1, and basal-like breast cancers with high expression of ALDH1 had higher expression of GLO1 and were more aggressive than cancers with low expression of ALDH1. Inhibition of GLO1 using TLSC702 in ALDH1-high cells induced cancer cell death and reduced tumor sphere formation, suggesting that GLO1 is a potential target for cancer therapy in terms of CSC survival [15]. Understanding the intricate role of Glo1 in cancer may provide valuable insights for the development of targeted therapies and personalized treatment strategies, ultimately impacting the clinical management and prognosis of cancer patients [16]. In this study, we identified the upregulation of GLO1 in the tissues of cervical cancer patients and validated the antitumor effects of a proven GLO1 inhibitor (BBGC) in cervical cancer cell lines. Thus, we demonstrate the potential of GLO1 as a promising therapeutic target for cervical cancer and suggest that application of GLO1 inhibitors as adjuvant chemotherapy in cancers with high GLO1 expression may increase therapeutic efficacy and improve survival.

## Materials and Methods

### Immunohistochemistry

Cervical cancer patient samples were obtained with approval from the Institutional Review Board (IRB) of Kangwon National University Hospital (KNUH-A-2022-07-010). The biospecimens and data used for this study were provided by the Biobank of Kangwon University Hospital, a member of the Korea Biobank Network.

Tissues stored in the Biobank, along with anonymized participant information, are all preserved with prior consent from the patients. The collection dates for the participant tissues span from January 1, 2021, to December 31, 2022. Information regarding individual medical records of participants is inaccessible, and only anonymized patient information can be obtained from the Biobank. All participants providing specimens for storage in the Biobank have given prior consent, which has been documented in written form and is held by the Biobank. Minors are not included among the participants.

Histological data, including patient age, gender, stage, and pathology, are summarized in Table 1. Paraffin-embedded tissue sections were deparaffinized and rehydrated. The slides underwent antigen retrieval using a buffer (0.01 M sodium citrate, pH 6) and were quenched in 3% H_2_O_2_ for 15 min. For GLO1 staining, sections were incubated with GLO1 antibody (ab171121) overnight at 4 °C. The labeled antigen was visualized using DAKO secondary antibody (DAKO) and DAB solution (ab64238; Abcam Inc, Toronto, ON, Canada). Sections were counterstained with hematoxylin and then mounted. The intensity of GLO1 protein staining in the tissues was evaluated on a scale of 0 to 5, with 0 indicating no staining (negative), 1 for weak staining, 2 to 3 for moderate staining, and 4 to 5 for strong staining. Low expression was defined as negative or weak staining, and high expression as moderate or strong staining. Each section was independently evaluated by three researchers.

**Table 1.**
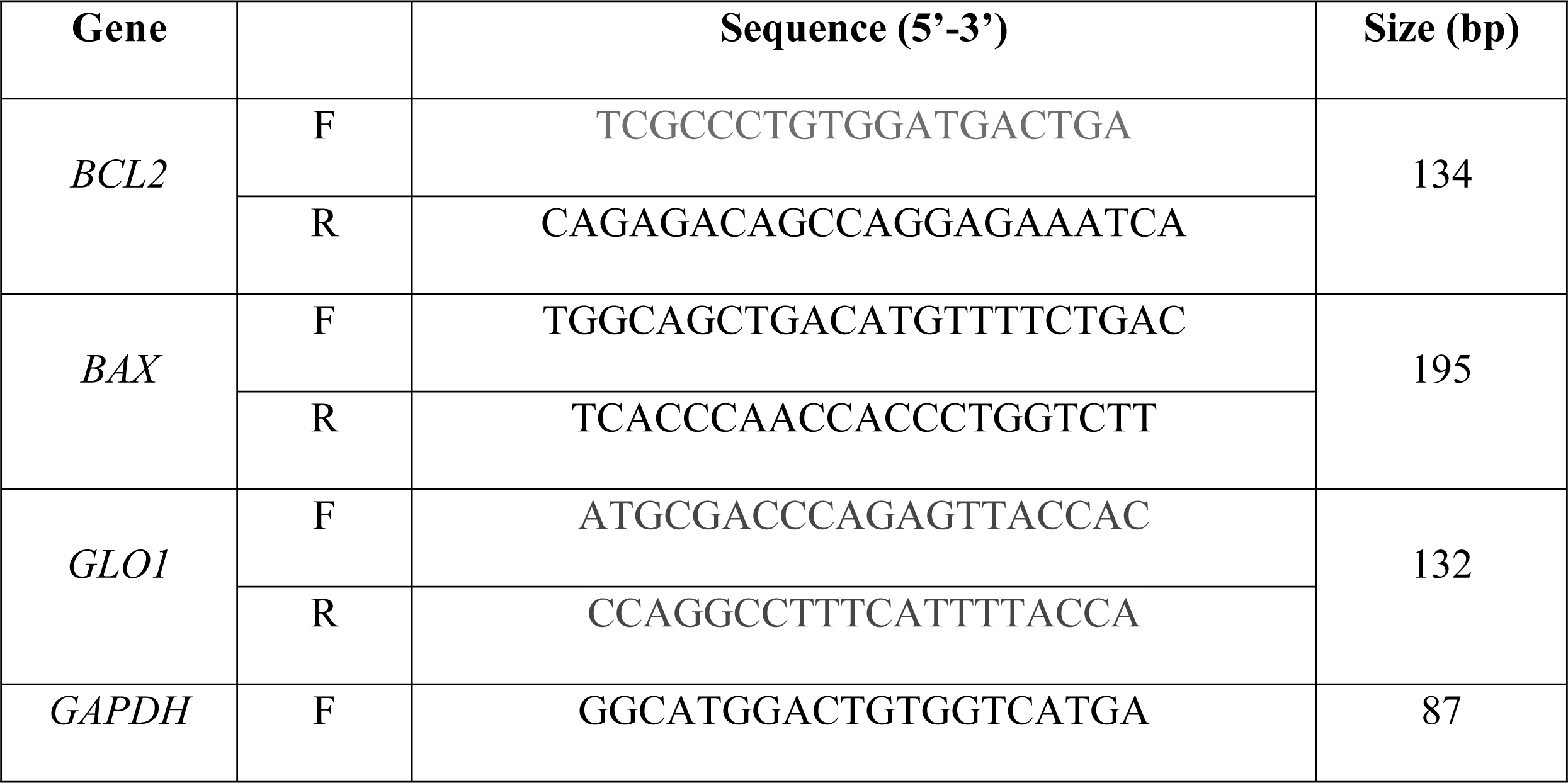
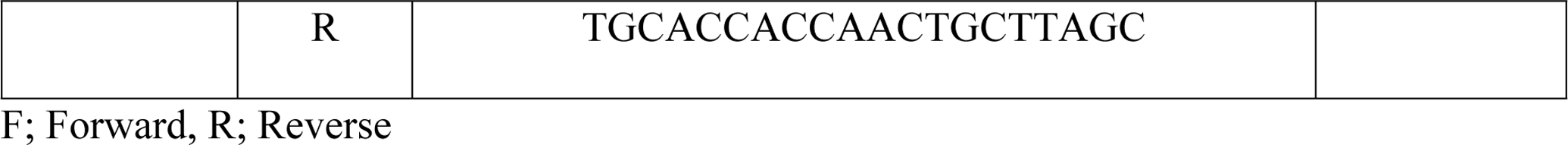
Primer sequences for real-time RT-PCR.

### Cell lines

HeLa cell lines were cultured in DMEM/High glucose (Thermo Fisher Scientific, Inc., Waltham, MA, USA), supplemented with 10% fetal bovine serum (FBS) (Thermo Fisher Scientific, Inc., Waltham, MA, USA) and 1% penicillin-streptomycin (Sigma-Aldrich, St. Louis, MO, USA). The cultures were maintained in a humidified atmosphere at 37 °C with 5% CO_2_. Upon reaching 80–90% confluency, the cells were seeded in 96-well plates for the MTT assay.

### MTT assay

24 hours post seeding of the HeLa cell line into 96-well plates, cells were treated with GLO1 inhibitor (SML1306; Sigma-Aldrich, St. Louis, MO, USA) at concentrations of 0 (control), 8 μM, and 10 μM for 48 h. After treatment, the cultures were replaced with serum-free medium containing MTT reagent (ab211091; Abcam Inc, Toronto, ON, Canada) and incubated for 3 h. The cells were subsequently treated with MTT solvent for 15 min at room temperature. Absorbance at 570 nm was measured using a microplate reader.

### Wound healing assay

2×10^5^ HeLa cells were seeded into 6-well plate culture dishes. The cells were treated with 0 (control), 8 μM, and 10 μM of GLO1 inhibitor for 24 h. For the wound healing assay, lines were created by scraping the cell monolayer with a 1000 μl pipette tip. Photos were taken under a microscope at 24, 48, and 72 h to document the closure of the wound.

### RNA extraction and quantitative real time PCR

Total RNA was extracted from HeLa cells using an RNeasy Mini kit (Qiagen, Duesseldorf, Germany) and cDNA was synthesized using TOPscrip^TM^ RT DryMIX (Enzynomics, Daejeon, Korea). PCR amplification was performed using a Step One Plus real time PCR system (Applied Biosystems, Warrington, UK) with TOPreal^TM^ qPCR 2X PreMIX (Enzynomics). The mRNA expression was normalized to an internal control GAPDH. The primer sequences are listed in Table 1.

### Single-cell RNA sequencing (scRNA-seq) data analysis

The scRNA-seq data from tumor and adjacent normal samples of cervical squamous cell carcinoma patients were downloaded from ArrayExpress (accession number E-MTAB-11948) [17]. We processed the scRNA-seq data using Seurat (v4.1.1) [18], focusing on gene-count matrices. For quality control, cells with a gene count ≤ 200 or a percentage of mitochondrial genes ≥ 20% were excluded as low-quality cells. The QC-passed data were log-normalized using the “NormalizeData” function. Next, we scaled the data by regressing out the percentage of mitochondrial genes and cell cycle scores with the “ScaleData” function. The cell cycle scores (S and G2M scores) were calculated using the “CellCycleScoring” function, based on the expression levels of S/G2M phase markers from Seurat. Principal component analysis (PCA) was performed on the scaled data using the top 2000 variable genes. For batch correction and sample integration, we utilized Harmony (v0.1.1) [19]. Unsupervised clustering was executed using “FindNeighbors” and “FindClusters” functions, with the results visualized via the “RunUMAP” function. Prior to further analysis, Scrublet (v0.2.3) [20] was employed to identify and remove a doublet cluster (**Fig. S1A and B**). Cell types for each cluster were annotated based on canonical marker genes and the top-ranked differentially expressed genes (**Fig. S1C**).

### Gene set variation analysis (GSVA)

GSVA (v1.42.0) [21] was employed to calculate the gene set enrichment scores for each cell. We downloaded Wikipathways gene sets (v2023.1) [22] from MSigDB [23, 24] to serve as the reference gene sets. The GSVA scores were then utilized in limma (v3.50.3) [25] to identify gene sets differentially regulated between normal and tumor epithelial cells. Pathways with an adjusted *p-value* < 0.01 were considered significant.

### Microarray data analysis

The microarray data for normal, cervical intraepithelial neoplasia (CIN1 to CIN3), and cervical cancer tissue samples were downloaded from the Gene Expression Omnibus (accession number GSE63514) [26]. Raw expression values were normalized using the Robust Multi-array Average (RMA) method with the affy package in R (version 1.72.0) [27].

### Statistical analysis

Values representing the expression of GLO1 in cervical cancer are expressed as means ± standard error of the mean (SEM). Statistical comparisons between groups were conducted using the *t*-test in GraphPad Prism software (GraphPad Software Inc, San Diego, CA, USA). Significance levels were set at *p* < 0.05 and *p* < 0.01.

## Results

### Populations

This study included a total of 58 patients diagnosed with cervical cancer, and their specimens were analyzed (**Table 2**). The average age of these patients was 55.1 years, with a standard deviation of ±15.08. Regarding cancer stages, 18 patients (31.03%) were in stage I, 16 patients (27.59%) in stage II, 8 patients (13.79%) in stage III, and 16 patients (27.59%) in stage IV. The histological analysis showed two primary types: squamous cell carcinoma in 52 patients (89.66%) and adenocarcinoma in 6 patients (10.34%). Immunohistochemistry specifically for glyoxalase 1 was conducted on specimens from all participating patients.

**Table 2.** Clinical Characteristics of Patients with Cervical Cancer.

| Variables | Cervical Cancer (n=58) | Control (n=16) |
| --- | --- | --- |
| Age (years) | 55.1±15.1 | 49.7±8.0 |
| Stage |  |  |
| I | 18 (31.03%) |  |
| II | 16 (27.59%) |  |
| III | 8 (13.79%) |  |
| IV | 16 (27.59%) |  |
| Histology |  |  |
| Squamous Cell Carcinoma | 52 (89.66%) |  |
| Adenocarcinoma | 6 (10.34%) |  |

Note: Data available for 58 patients with cervical cancer. SD, standard deviation.

### GLO1 is upregulated in cervical cancer tissues compared to normal cervical tissues

To compare the expression analysis of Glo1 in cervical cancer, tissue slides were obtained from cervix and ovary and Glo1 staining was performed. In normal tissues, the expression of GLO1 tends to be lower in cervical tissues compared to endometrium and ovary normal tissues, and in cancer tissues, the expression of GLO1 is upregulated in cervical cancer tissues. In particular, the expression of GLO1 is higher in cervical cancer tissues compared to normal, and the expression of GLO1 in endometrium and ovarian cancer tissues is not significantly different from normal and cancer tissues(Fig1A). Moreover, the staining intensity by IHC score was 3.372±0.340 in cervical cancer tissues compared to 0.260±0.164 in normal (Fig1B). These results show that the expression of GLO1 is specifically upregulated in cervical cancer tissues, supporting the potential involvement of GLO1 in tumor progression in the cervix.

**Fig 1.**
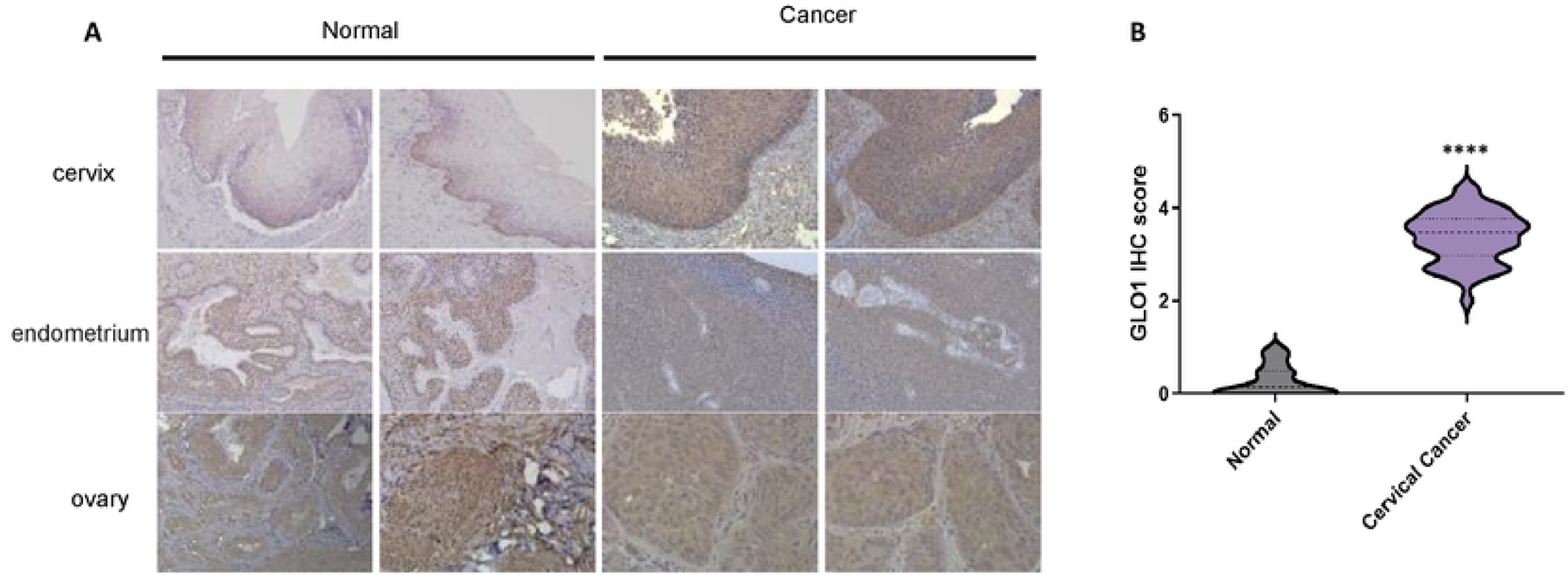
Upregulation of GLO1 in Cervical Cancer Tissues. (A) Representative staining of GLO1 in cervical cancer and normal tissues (200x magnification). (B) Comparative GLO1 IHC scores in cervical cancer versus normal tissues. Statistical significance was determined using the *t*-test (**** *p*<0.0001).

### Inhibiting GLO1 affects the proliferation and apoptosis of cervical cancer cells

To support the clinical findings that upregulation of Glo1 is involved in tumor progression, we inhibited the activity of Glo1 using BBGC, a Glo1 inhibitor, and evaluated the proliferation and apoptosis of cervical cancer cell lines. In Hela cells treated with 8 μM and 10 μM of BBGC, we observed the wound closure rate for 72 hours using the wound healing assay (Figure 2A). BBGC reduced the proliferation of cervical cancer cell lines in a dose- and time-dependent manner (Figure 2B), and to evaluate the effect of GLO1 downregulation on the viability of Hela cells, cell viability was measured by MTT assay. We found that the viability of the cells decreased with increasing concentrations of BBGC (Figure 2C), and the expression of BAX2, a pro-apoptosis-related gene, increased (Figure 2D). These results support that upregulation of Glo1 promotes the survival of cervical cancer cells and induces tumor metastasis, and further suggest that modulation of Glo1 may be a target for anticancer chemotherapy.

**Fig 2.**
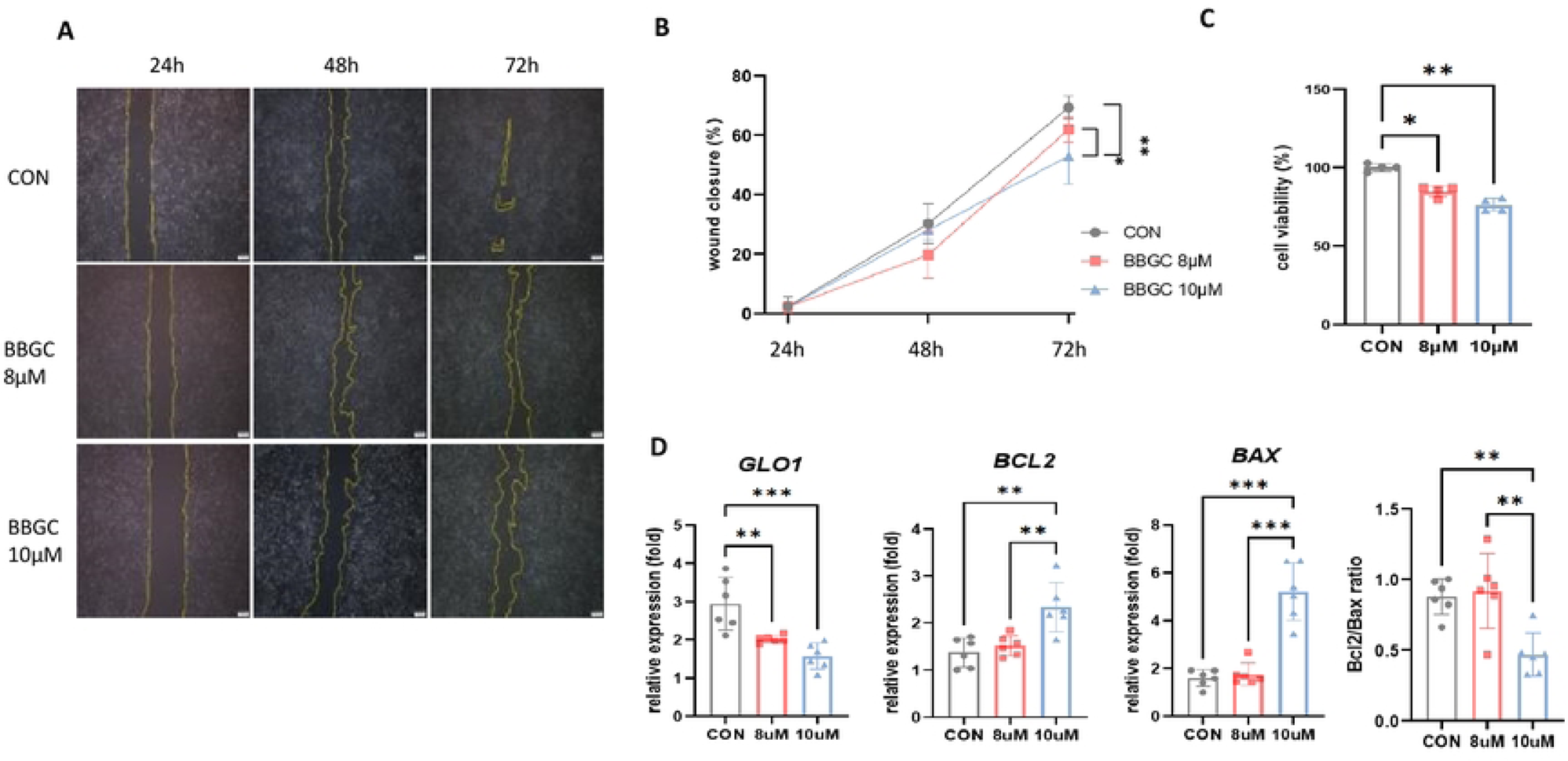
Impact of GLO1 Inhibitor on HeLa Cell Viability and Migration. (A) Wound healing assay evaluating the effect of GLO1 inhibitor (BBGC) on HeLa cell migration at 24, 48, and 72 h. (B) Quantitative analysis of HeLa cell migration distance. (C) Assessment of HeLa cell viability with 8 μM and 10 μM GLO1 inhibitor using the MTT assay. Statistical significance was analyzed using the *t*-test (*** *p*<0.001; ** *p*<0.01; * *p*<0.05).

### scRNA-seq analysis of adjacent normal and tumor samples from cervical squamous cell carcinoma patients (E-MTAB-11948)

To corroborate our experimental findings, we delved into public gene expression data for cervical cancer at both single-cell and bulk levels. We discovered scRNA-seq data for tumor-normal paired tissues from two patients (ArrayExpress E-MTAB-11948). After performing quality control, normalization, and integration, we successfully identified epithelial, stromal, and various immune cell types (**Fig. 3A**). Epithelial cells were notably predominant in the tumor tissue (**Fig. 3B**). *GLO1* expression was found to be significantly higher in tumor epithelial cells compared to normal epithelial cells (**Fig. 3C**). We then analyzed which pathways were differentially regulated between normal and tumor epithelial cells. Cell-specific pathway activities were determined using the GSVA algorithm for Wikipathways, revealing that many pathways enriched in tumor cells were associated with the glycolysis process (**Fig. 3D**). Overall, the single-cell data confirmed that *GLO1* is overexpressed in cervical tumor samples, potentially playing a crucial role in cancer metabolism through the glycolysis pathway.

**Fig 3.**
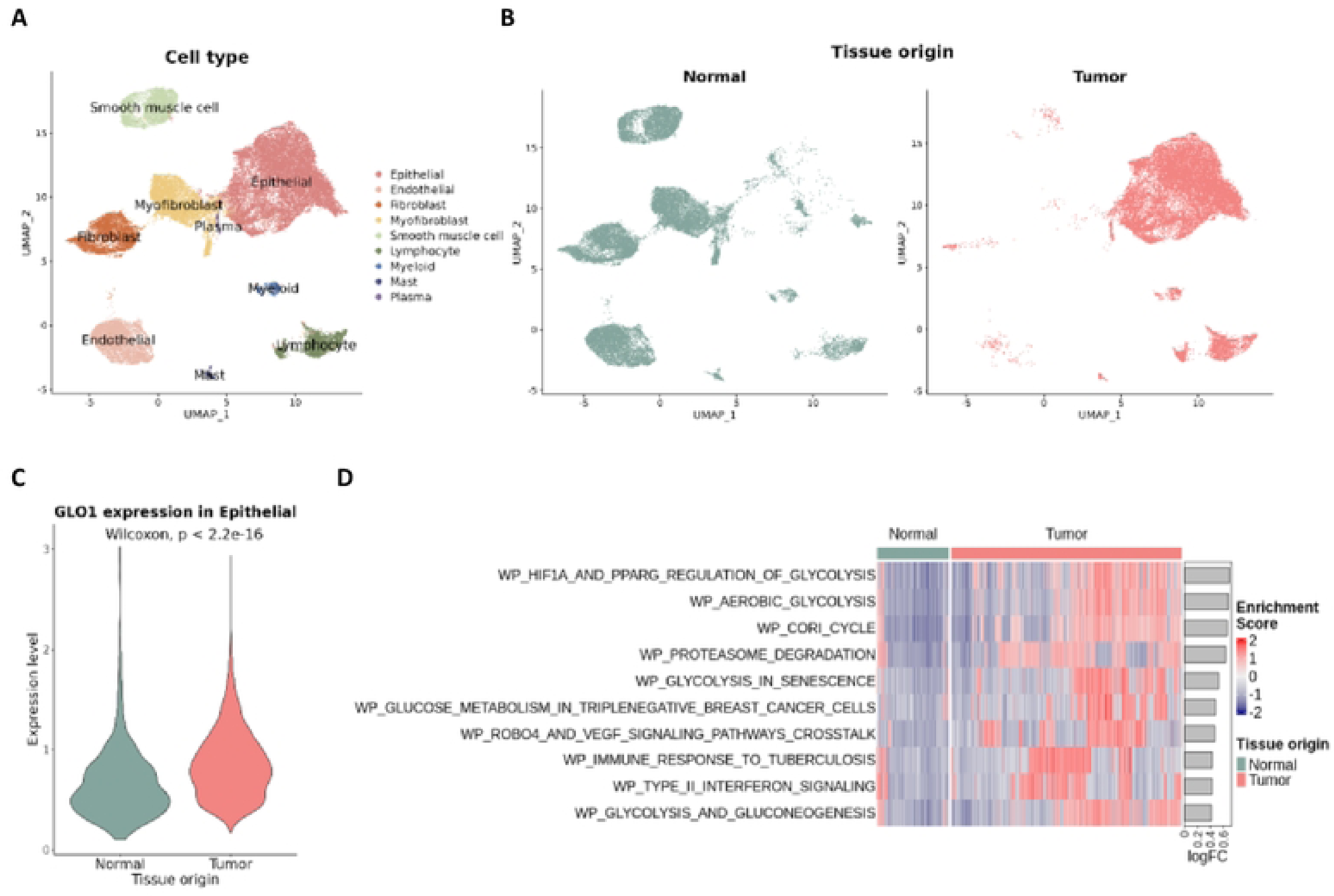
scRNA-seq Analysis of *GLO1* Expression in Cervical Squamous Cell Carcinoma. (A) UMAP plot of 44,848 cells categorized by cell types. (B) UMAP distribution of 23,636 normal and 21,212 tumor cells. (C) *GLO1* expression in epithelial cells from normal and tumor tissues, excluding cells not expressing GLO1. (D) Heatmap of GSVA enrichment scores for Wikipathways gene sets, comparing epithelial cells from normal and tumor tissues. The top 10 pathways are displayed based on the logFC of average GSVA scores between the two groups.

### *GLO1* Expression in normal, cervical intraepithelial neoplasia (CIN1 to CIN3), and cervical cancer tissue samples

To investigate whether GLO1 expression correlates with cancer progression, we analyzed public expression data of cervical cancer samples across various tumor stages. The GEO GSE63514 microarray data set included 24 normal, 14 cervical intraepithelial neoplasia (CIN) 1, 22 CIN2, 40 CIN3, and 28 cervical cancer samples. We observed that *GLO1* expression progressively increased from the normal to CIN1, CIN2, CIN3, and tumor sample groups (**Fig. 4**). Our analysis independently confirms that *GLO1* is overexpressed in cervical tumors, likely in a progression-dependent manner.

**Fig 4.**
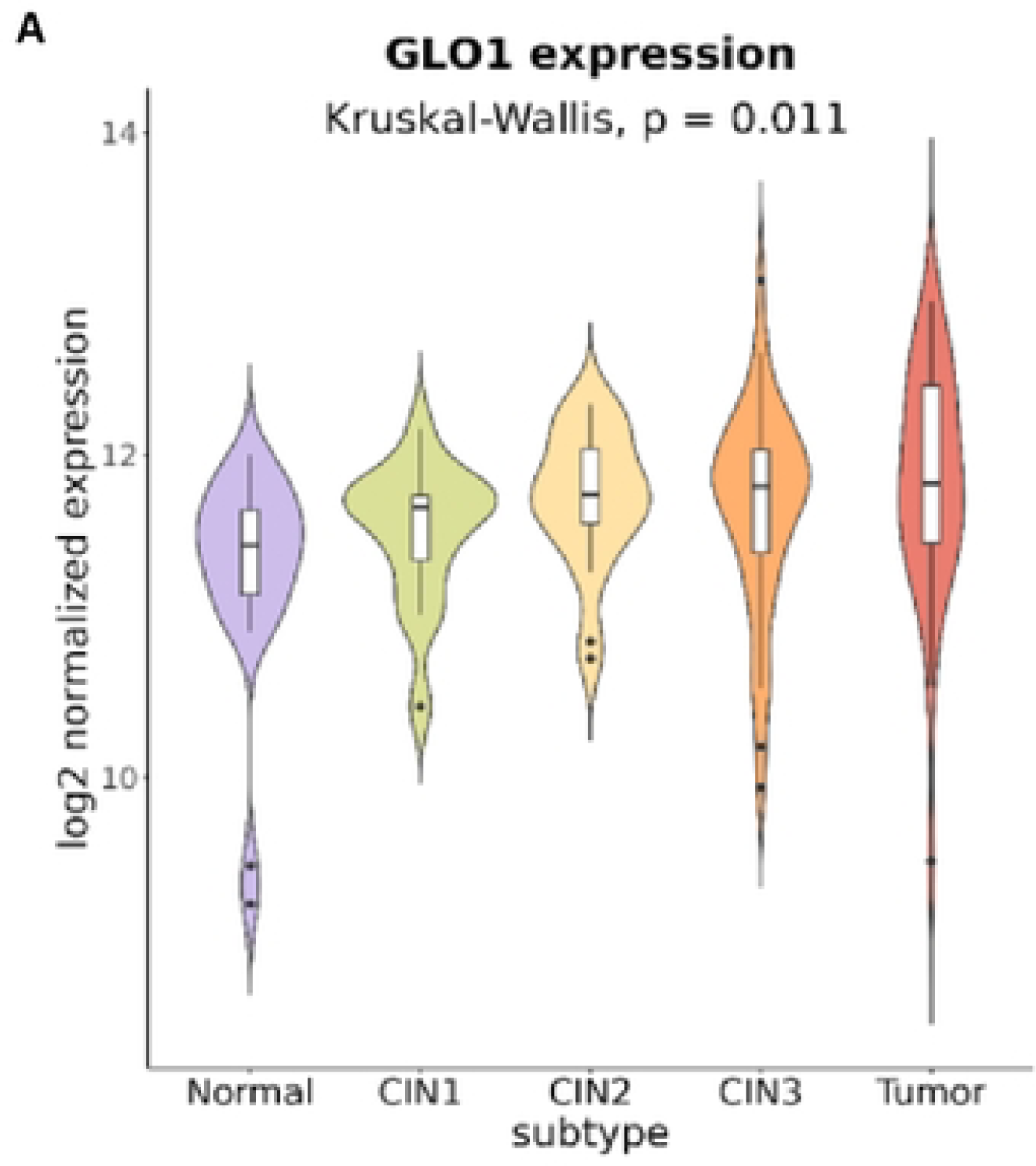
GLO1 Expression in Cervical Tissue Progression Stages. Microarray analysis of *GLO1* expression in normal, cervical intraepithelial neoplasia (CIN1 to CIN3), and cervical cancer tissues. Data from GEO database (GSE63514) visualized in violin plots for 24 normal, 14 CIN1, 22 CIN2, 40 CIN3, and 28 cervical cancer samples.

## Discussion

In this study, we examined 58 cervical cancer patients, analyzing their tissue specimens. We found that Glyoxalase 1 (GLO1) is significantly upregulated in cervical cancer tissues compared to normal cervical tissues, irrespective of the pathological findings and stage. Intriguingly, our findings align with proteomic profiling data from the Human Protein Atlas (v23.proteinatlas.org), which ranks GLO1 among the top 10 most critical proteins in cancer prediction models, scoring from 0 to 100 (Fig. S2A). In pan-cancer protein panel analysis for predicting cervical cancer, GLO1 emerges as the most significant among the seven key proteins (Fig. S2B). This upregulation of GLO1 correlates with increased cell viability and migration in cervical cancer cell lines, underscoring its contribution to tumor progression and its potential as a chemotherapy target. Additionally, single-cell analysis and scrutiny of public gene expression data have confirmed GLO1’s overexpression in cervical tumor samples and its role in cancer metabolism, particularly in the glycolysis pathway. Moreover, the evaluation of public expression data from precancerous lesions, including CI 1 to 3, indicates that *GLO1* expression progressively increased from normal to precancerous lesions and invasive cervical cancer, suggesting a progression-dependent association with carcinogenesis.

GLO1 is a zinc-dependent metalloenzyme comprised of two identical subunits, each with a molecular mass of 43–48 kDa [28]. It is ubiquitously present in both prokaryotic and eukaryotic organisms, and its high homology across different species suggests a function that is evolutionarily conserved. In humans, *GLO1* is located on chromosome 6 and is often overexpressed in tumor tissues [29]. The regulation of *GLO1* expression is multifaceted, involving various regulatory elements, gene expression alterations, and post-translational modifications such as phosphorylation, nitrosylation, and glutathionylation [30]. Transcriptional regulators of GLO1 encompass AP-2α, E2F4, NF-κB, AP-1, ARE, MRE, and IRE elements [31]. Post-translational modifications occur through phosphorylation, nitrosylation, and glutathionylation. Nrf2 and its activators contribute to the upregulation of GLO1 expression under stress conditions [30]. Conversely, GLO1 expression is negatively regulated by HIF1α and RAGE [32]. Copy number variations (CNVs) of the *GLO1* gene in the human genome can result in increased *GLO1* expression [33], a phenomenon more prevalent in various human tumors, including breast cancer, sarcomas, and non-small cell lung cancer [34].

Cancer metabolism is marked by the reprogramming of cellular energy production and nutrient utilization, particularly through heightened glycolysis, even in oxygen-rich environments, a phenomenon known as the Warburg effect [35, 36]. In cervical cancer, this metabolic shift is crucial for tumor progression [37]. The enhanced glycolysis in cervical cancer cells, fueled by factors such as GLO1 overexpression, not only generates energy and biosynthetic intermediates for rapidly proliferating cancer cells but also contributes to the formation of an acidic tumor microenvironment and chemoresistance. This ultimately supports cancer cell survival and invasiveness [34]. Understanding these specific metabolic changes in cervical cancer, like glycolysis dysregulation and the role of key enzymes such as GLO1, presents valuable opportunities for targeted therapies and the development of new diagnostic and prognostic tools in combating this widespread and potentially lethal disease.

The association between HPV infection and the progression of cervical cancer, particularly from a metabolic perspective, is significant. Human papillomavirus (HPV) infection, especially with high-risk HPV types, is a recognized risk factor for developing cervical cancer [38]. The process initiates when the virus infects cervical epithelial cells, and its oncoproteins, E6 and E7, disrupt key cellular regulatory pathways [39]. The elevated levels of GLO1 may contribute to the evasion of apoptosis and the enhanced survival of HPV-infected cells [40]. Additionally, the glyoxalase system, which includes GLO1, plays a role in controlling cellular reactive oxygen species (ROS) levels, and the dysregulation of ROS is a notable characteristic in cancer development [41]. From a metabolic perspective, these viral oncoproteins interfere with the functions of tumor suppressor proteins, such as p53 and Rb, inducing significant changes in cellular metabolism [42]. This metabolic alteration enables infected cells to favor glycolysis for energy production, even in oxygen-rich environments. This shift not only supports the rapid proliferation of cancer cells but also contributes to an acidic tumor microenvironment, potentially enhancing cancer cell invasiveness [43]. Moreover, the disruption of glucose metabolism leads to alterations in various cellular pathways, including those critical for the synthesis of nucleic acids, lipids, and amino acids, essential for cancer cell growth and division [44]. These metabolic adaptations are key to the progression of cervical cancer, from the early stages of HPV infection to the development of precancerous lesions and ultimately invasive cancer.

In clinical practice, no tissue biological markers are currently used to refine prognosis or customize treatment strategies for cervical carcinoma patients. However, a thorough review of the literature reveals that the presence of the HPV-18 genotype and elevated VEGF expression correlate with poorer prognoses in women with early-stage disease who undergo surgical treatment [45, 46]. Conversely, the expression of EGFR, VEGF, COX-2, and tumor hypoxia significantly influences the survival of patients receiving definitive radiotherapy or chemoradiation [47]. Although the evaluation of targeted therapies in cervical carcinoma is limited and lacks published phase III trials or approved agents in clinical practice, it is notable that high-risk HPVs can promote the production of angiogenic factors, likely through the upregulation of E6 and E7 oncoproteins [48]. In HPV-16 positive cells, the expression of pro-angiogenic molecules such as βFGF, IL-8, TGF-beta, and VEGF is higher compared to control keratinocytes [49, 50]. The adverse prognostic impact of increased tumoral VEGF expression and the encouraging outcomes of bevacizumab in therapeutic trials for cervical carcinoma indicate that targeting angiogenesis pathways is a promising therapeutic strategy for this disease. The identification of new cancer metabolism markers like Glo-1 will enhance our understanding of cervical cancer in the future, underscoring the need for ongoing research.

There were several limitations to our study. Firstly, the sample size was relatively small, preventing us from identifying significant differences in GLO1 expression across different stages of cervical cancer (early versus advanced or metastatic). Nevertheless, it is noteworthy that we have identified a meaningful role for GLO1 in cervical cancer for the first time. Secondly, we did not explore GLO1’s potential as a biomarker in other biological samples, such as patient serum or urine, which warrants further investigation. Despite these limitations, our study possesses several strengths. Primarily, it contributes to the identification of a novel biomarker, GLO1, in cervical cancer. Our research highlights the significant upregulation of *GLO1* in cervical cancer tissues, supporting its potential as a valuable biomarker for tumor progression and as a target for cancer therapy. Moreover, we employed a comprehensive methodology that integrates clinical analysis of patient samples, *in vitro* experiments with cell lines, single-cell RNA sequencing (scRNA-seq) data analysis, and examination of public gene expression data. This multifaceted approach offers a thorough understanding of GLO1’s role in cervical cancer, thereby strengthening the credibility of our findings.

## Conclusion

Overall, our comprehensive analysis underscores the pivotal role of GLO1 in the progression of cervical cancer, providing fresh perspectives on its viability as both a therapeutic target and biomarker for this life-threatening disease. Grasping the molecular intricacies of cervical cancer is essential for devising novel diagnostic and treatment approaches. Our findings make a significant contribution to the current endeavors aimed at combating cervical cancer.

## Conflict of Interest

No potential conflict of interest relevant to this article was reported.

## Data sharing statement

The datasets utilized and/or analyzed during this study are available from the corresponding author upon reasonable request.

## Author Contributions

Conceptualization: H.S.H., O.Y.T.; Data Curation: K.J.Y., J.J.H., O.Y.T., L.H.A.; Formal Analysis: J.S.R., L.S.H.; Investigation: H.S.H., O.Y.T., K.J.Y., J.J.H., L.H.A.; Methodology: K.J.Y.; Resources: O.Y.T.; Supervision: H.S.H.; Visualization: K.J.Y., J.J.H., J.S.R., L.S.H.; Writing - Original Draft: O.Y.T., K.J.Y.; Writing - Review & Editing: H.S.H., O.Y.T.

## Fundings

This study was supported by the National Research Foundation of Korea (NRF) grant funded by Korea (RS-2023-00279214).

## Acknowledgement

This study was also supported by the Analytic Biological Service Project (ABSP), funded by the Kangwon National University Hospital Grant in 2021.

This study was supported by the Analytic Biological Service Project (ABSP) funded by the Kangwon National University Hospital Grant in 2022.

The biospecimens and data used for this study were provided by the Biobank of Kangwon University Hospital, a member of the Korea Biobank Network.

We would like to thank Editage (www.editage.co.kr) for English language editing.

## Notes

### Competing Interest Statement

The authors have declared no competing interest.

## References

1. Ginsburg O, Bray F, Coleman MP, Vanderpuye V, Eniu A, Kotha SR, et al. The global burden of women’s cancers: a grand challenge in global health. Lancet. 2017;389(10071):847–60. 10.1016/s0140-6736(16)31392-7 PMID: 27814965

2. Cohen CM, Wentzensen N, Castle PE, Schiffman M, Zuna R, Arend RC, Clarke MA. Racial and Ethnic Disparities in Cervical Cancer Incidence, Survival, and Mortality by Histologic Subtype. J Clin Oncol. 2023;41(5):1059–68. 10.1200/jco.22.01424 PMID: 36455190

3. Sokale IO, Oluyomi AO, Montealegre JR, Thrift AP. Racial/Ethnic Disparities in Cervical Cancer Stage at Diagnosis: Mediating Effects of Neighborhood-level Socioeconomic Deprivation. Cancer Epidemiol Biomarkers Prev. 2023;32(6):818–24. 10.1158/1055-9965.Epi-23-0038 PMID: 37067295

4. Trunk MJ, Wentzensen N, von Knebel Doeberitz M. [Molecular pathogenesis of cervical cancer and its first steps]. Pathologe. 2005;26(4):283–90. 10.1007/s00292-005-0763-4 PMID: 15928953

5. Balasubramaniam SD, Balakrishnan V, Oon CE, Kaur G. Key Molecular Events in Cervical Cancer Development. Medicina (Kaunas). 2019;55(7). 10.3390/medicina55070384 PMID: 31319555

6. Kidd EA, Grigsby PW. Intratumoral metabolic heterogeneity of cervical cancer. Clin Cancer Res. 2008;14(16):5236–41. 10.1158/1078-0432.Ccr-07-5252 PMID: 18698042

7. Antognelli C, Cecchetti R, Riuzzi F, Peirce MJ, Talesa VN. Glyoxalase 1 sustains the metastatic phenotype of prostate cancer cells via EMT control. J Cell Mol Med. 2018;22(5):2865–83. 10.1111/jcmm.13581 PMID: 29504694

8. Antognelli C, Mezzasoma L, Mearini E, Talesa VN. Glyoxalase 1-419C>A variant is associated with oxidative stress: implications in prostate cancer progression. PLoS One. 2013;8(9):e74014. 10.1371/journal.pone.0074014 PMID: 24040147

9. Baunacke M, Horn LC, Trettner S, Engel KM, Hemdan NY, Wiechmann V, et al. Exploring glyoxalase 1 expression in prostate cancer tissues: targeting the enzyme by ethyl pyruvate defangs some malignancy-associated properties. Prostate. 2014;74(1):48–60. 10.1002/pros.22728 PMID: 24105621

10. Motomura H, Ozaki A, Tamori S, Onaga C, Nozaki Y, Waki Y, et al. Glyoxalase 1 and protein kinase Cλ as potential therapeutic targets for late-stage breast cancer. Oncol Lett. 2021;22(1):547. 10.3892/ol.2021.12808 PMID: 34093768

11. Tamori S, Nozaki Y, Motomura H, Nakane H, Katayama R, Onaga C, et al. Glyoxalase 1 gene is highly expressed in basal-like human breast cancers and contributes to survival of ALDH1-positive breast cancer stem cells. Oncotarget. 2018;9(92):36515–29. 10.18632/oncotarget.26369 PMID: 30559934

12. Zhang Z, Dong L, Sui L, Yang Y, Liu X, Yu Y, et al. Metformin reverses progestin resistance in endometrial cancer cells by downregulating GloI expression. Int J Gynecol Cancer. 2011;21(2):213–21. 10.1097/IGC.0b013e318207dac7 PMID: 21270604

13. Alhujaily M, Abbas H, Xue M, de la Fuente A, Rabbani N, Thornalley PJ. Studies of Glyoxalase 1-Linked Multidrug Resistance Reveal Glycolysis-Derived Reactive Metabolite, Methylglyoxal, Is a Common Contributor in Cancer Chemotherapy Targeting the Spliceosome. Front Oncol. 2021;11:748698. 10.3389/fonc.2021.748698 PMID: 34790575

14. Thornalley PJ. Protecting the genome: defence against nucleotide glycation and emerging role of glyoxalase I overexpression in multidrug resistance in cancer chemotherapy. Biochem Soc Trans. 2003;31(Pt 6):1372–7. 10.1042/bst0311372 PMID: 14641066

15. Ricardo S, Vieira AF, Gerhard R, Leitão D, Pinto R, Cameselle-Teijeiro JF, et al. Breast cancer stem cell markers CD44, CD24 and ALDH1: expression distribution within intrinsic molecular subtype. J Clin Pathol. 2011;64(11):937–46. 10.1136/jcp.2011.090456 PMID: 21680574

16. Geng X, Ma J, Zhang F, Xu C. Glyoxalase I in tumor cell proliferation and survival and as a potential target for anticancer therapy. Oncol Res Treat. 2014;37(10):570–4. 10.1159/000367800 PMID: 25342507

17. Li C, Wu H, Guo L, Liu D, Yang S, Li S, Hua K. Single-cell transcriptomics reveals cellular heterogeneity and molecular stratification of cervical cancer. Commun Biol. 2022;5(1):1208. 10.1038/s42003-022-04142-w PMID: 36357663

18. Hao Y, Hao S, Andersen-Nissen E, Mauck WM, 3rd, Zheng S, Butler A, et al. Integrated analysis of multimodal single-cell data. Cell. 2021;184(13):3573–87.e29. 10.1016/j.cell.2021.04.048 PMID: 34062119

19. Korsunsky I, Millard N, Fan J, Slowikowski K, Zhang F, Wei K, et al. Fast, sensitive and accurate integration of single-cell data with Harmony. Nat Methods. 2019;16(12):1289–96. 10.1038/s41592-019-0619-0 PMID: 31740819

20. Wolock SL, Lopez R, Klein AM. Scrublet: Computational Identification of Cell Doublets in Single-Cell Transcriptomic Data. Cell Syst. 2019;8(4):281–91.e9. 10.1016/j.cels.2018.11.005 PMID: 30954476

21. Hänzelmann S, Castelo R, Guinney J. GSVA: gene set variation analysis for microarray and RNA-seq data. BMC Bioinformatics. 2013;14:7. 10.1186/1471-2105-14-7 PMID: 23323831

22. Pico AR, Kelder T, van Iersel MP, Hanspers K, Conklin BR, Evelo C. WikiPathways: pathway editing for the people. PLoS Biol. 2008;6(7):e184. 10.1371/journal.pbio.0060184 PMID: 18651794

23. Subramanian A, Tamayo P, Mootha VK, Mukherjee S, Ebert BL, Gillette MA, et al. Gene set enrichment analysis: a knowledge-based approach for interpreting genome-wide expression profiles. Proc Natl Acad Sci U S A. 2005;102(43):15545–50. 10.1073/pnas.0506580102 PMID: 16199517

24. Liberzon A, Birger C, Thorvaldsdóttir H, Ghandi M, Mesirov JP, Tamayo P. The Molecular Signatures Database (MSigDB) hallmark gene set collection. Cell Syst. 2015;1(6):417–25. 10.1016/j.cels.2015.12.004 PMID: 26771021

25. Ritchie ME, Phipson B, Wu D, Hu Y, Law CW, Shi W, Smyth GK. limma powers differential expression analyses for RNA-sequencing and microarray studies. Nucleic Acids Res. 2015;43(7):e47. 10.1093/nar/gkv007 PMID: 25605792

26. den Boon JA, Pyeon D, Wang SS, Horswill M, Schiffman M, Sherman M, et al. Molecular transitions from papillomavirus infection to cervical precancer and cancer: Role of stromal estrogen receptor signaling. Proc Natl Acad Sci U S A. 2015;112(25):E3255–64. 10.1073/pnas.1509322112 PMID: 26056290

27. Gautier L, Cope L, Bolstad BM, Irizarry RA. affy--analysis of Affymetrix GeneChip data at the probe level. Bioinformatics. 2004;20(3):307–15. 10.1093/bioinformatics/btg405 PMID: 14960456

28. Cameron AD, Olin B, Ridderström M, Mannervik B, Jones TA. Crystal structure of human glyoxalase I--evidence for gene duplication and 3D domain swapping. Embo j. 1997;16(12):3386–95. 10.1093/emboj/16.12.3386 PMID: 9218781

29. Tripodis N, Mason R, Humphray SJ, Davies AF, Herberg JA, Trowsdale J, et al. Physical map of human 6p21.2-6p21.3: region flanking the centromeric end of the major histocompatibility complex. Genome Res. 1998;8(6):631–43. 10.1101/gr.8.6.631 PMID: 9647638

30. MacLeod AK, McMahon M, Plummer SM, Higgins LG, Penning TM, Igarashi K, Hayes JD. Characterization of the cancer chemopreventive NRF2-dependent gene battery in human keratinocytes: demonstration that the KEAP1-NRF2 pathway, and not the BACH1-NRF2 pathway, controls cytoprotection against electrophiles as well as redox-cycling compounds. Carcinogenesis. 2009;30(9):1571–80. 10.1093/carcin/bgp176 PMID: 19608619

31. Antognelli C, Palumbo I, Aristei C, Talesa VN. Glyoxalase I inhibition induces apoptosis in irradiated MCF-7 cells via a novel mechanism involving Hsp27, p53 and NF-κB. Br J Cancer. 2014;111(2):395–406. 10.1038/bjc.2014.280 PMID: 24918814

32. Aragonès G, Rowan S, Francisco SG, Whitcomb EA, Yang W, Perini-Villanueva G, et al. The Glyoxalase System in Age-Related Diseases: Nutritional Intervention as Anti-Ageing Strategy. Cells. 2021;10(8):1852. 10.3390/cells10081852 PMID: 34440621

33. Redon R, Ishikawa S, Fitch KR, Feuk L, Perry GH, Andrews TD, et al. Global variation in copy number in the human genome. Nature. 2006;444(7118):444-54. 10.1038/nature05329 PMID: 17122850

34. Kim JY, Jung JH, Lee SJ, Han SS, Hong SH. Glyoxalase 1 as a Therapeutic Target in Cancer and Cancer Stem Cells. Mol Cells. 2022;45(12):869–76. 10.14348/molcells.2022.0109 PMID: 36172978

35. Jang M, Kim SS, Lee J. Cancer cell metabolism: implications for therapeutic targets. Exp Mol Med. 2013;45(10):e45. 10.1038/emm.2013.85 PMID: 24091747

36. Ferreira LM. Cancer metabolism: the Warburg effect today. Exp Mol Pathol. 2010;89(3):372–80. 10.1016/j.yexmp.2010.08.006 PMID: 20804748

37. Li B, Sui L. Metabolic reprogramming in cervical cancer and metabolomics perspectives. Nutr Metab (Lond). 2021;18(1):93. 10.1186/s12986-021-00615-7 PMID: 34666780

38. So KA, Lee IH, Lee KH, Hong SR, Kim YJ, Seo HH, Kim TJ. Human papillomavirus genotype-specific risk in cervical carcinogenesis. J Gynecol Oncol. 2019;30(4):e52. 10.3802/jgo.2019.30.e52 PMID: 31074234

39. Tomaić V. Functional Roles of E6 and E7 Oncoproteins in HPV-Induced Malignancies at Diverse Anatomical Sites. Cancers (Basel). 2016;8(10). 10.3390/cancers8100095 PMID: 27775564

40. Yuan CH, Filippova M, Duerksen-Hughes P. Modulation of apoptotic pathways by human papillomaviruses (HPV): mechanisms and implications for therapy. Viruses. 2012;4(12):3831–50. 10.3390/v4123831 PMID: 23250450

41. He Y, Zhou C, Huang M, Tang C, Liu X, Yue Y, et al. Glyoxalase system: A systematic review of its biological activity, related-diseases, screening methods and small molecule regulators. Biomed Pharmacother. 2020;131:110663. 10.1016/j.biopha.2020.110663 PMID: 32858501

42. Pal A, Kundu R. Human Papillomavirus E6 and E7: The Cervical Cancer Hallmarks and Targets for Therapy. Front Microbiol. 2019;10:3116. 10.3389/fmicb.2019.03116 PMID: 32038557

43. Zhang Y, Yang JM. Altered energy metabolism in cancer: a unique opportunity for therapeutic intervention. Cancer Biol Ther. 2013;14(2):81–9. 10.4161/cbt.22958 PMID: 23192270

44. Schiliro C, Firestein BL. Mechanisms of Metabolic Reprogramming in Cancer Cells Supporting Enhanced Growth and Proliferation. Cells. 2021;10(5):1056. 10.3390/cells10051056 PMID: 33946927

45. Kang WD, Kim CH, Cho MK, Kim JW, Cho HY, Kim YH, et al. HPV-18 is a poor prognostic factor, unlike the HPV viral load, in patients with stage IB-IIA cervical cancer undergoing radical hysterectomy. Gynecol Oncol. 2011;121(3):546–50. 10.1016/j.ygyno.2011.01.015 PMID: 21334052

46. Ma DM, Xu YP, Zhu L. Expression of vascular endothelial growth factor C correlates with a poor prognosis based on analysis of prognostic factors in patients with cervical carcinomas. J Obstet Gynaecol Res. 2011;37(11):1519–24. 10.1111/j.1447-0756.2011.01566.x PMID: 21676077

47. Yoysungnoen-Chintana P, Bhattarakosol P, Patumraj S. Antitumor and antiangiogenic activities of curcumin in cervical cancer xenografts in nude mice. Biomed Res Int. 2014;2014:817972. 10.1155/2014/817972 PMID: 24860830

48. Fleischmann M, Chatzikonstantinou G, Fokas E, Wichmann J, Christiansen H, Strebhardt K, et al. Molecular Markers to Predict Prognosis and Treatment Response in Uterine Cervical Cancer. Cancers (Basel). 2021;13(22):5748. 10.3390/cancers13225748 PMID: 34830902

49. Marquina G, Manzano A, Casado A. Targeted Agents in Cervical Cancer: Beyond Bevacizumab. Curr Oncol Rep. 2018;20(5):40. 10.1007/s11912-018-0680-3 PMID: 29611060

50. Zagouri F, Sergentanis TN, Chrysikos D, Filipits M, Bartsch R. Molecularly targeted therapies in cervical cancer. A systematic review. Gynecol Oncol. 2012;126(2):291–303. 10.1016/j.ygyno.2012.04.007 PMID: 22504292

